# The influence of environment geometry on subiculum boundary vector cells in adulthood and early development

**DOI:** 10.1101/2023.04.13.536690

**Authors:** Laurenz Muessig, Fabio Ribeiro Rodrigues, Tale Bjerknes, Ben Towse, Caswell Barry, Neil Burgess, Edvard I. Moser, May-Britt Moser, Francesca Cacucci, Thomas J. Wills

## Abstract

Boundaries to movement form a specific class of landmark information used for navigation: Boundary Vector Cells (BVCs) are neurons which encode an animal’s location as a vector displacement from boundaries. Here we report the first objective characterisation of the prevalence and spatial tuning of subiculum BVCs. Manipulations of boundary geometry reveal two novel features of BVC firing. Firstly, BVC directional tunings align with environment walls in squares, but are uniformly distributed in circles, demonstrating that environmental geometry alters BVC receptive fields. Secondly, inserted barriers uncover both excitatory and inhibitory components to BVC receptive fields, demonstrating that inhibitory inputs contribute to BVC field formation. During post-natal development, subiculum BVCs mature slowly, contrasting with the earlier maturation of boundary-responsive cells in upstream Entorhinal Cortex. However, Subiculum and Entorhinal BVC receptive fields are altered by boundary geometry as early as tested, suggesting this is an inherent feature of the hippocampal representation of space.

## Introduction

Spatial cognition in the hippocampus is supported by a network of neurons tuned to an animal’s position and orientation, including place^1^, head direction^2^ and grid^3^ cells, which collectively form a cognitive map of allocentric space^4, 5^. The spatial tuning of these neurons is supported by a combination of both internally-derived movement information and sensory-bound external landmarks^6–8^. One important class of external landmarks are environmental boundaries, more specifically, extended objects that form a barrier to an animal’s movement^9, 10^. Boundaries are thought to anchor the cognitive map at the edges of the visited environment, correcting errors accumulated in an open field^11–16^.

At a behavioural level, boundary geometry serves as a strong cue for the successful retrieval of spatial memories. After disorientation, many species of animals, including human children, will search for a reward in geometrically equivalent corners of a rectangular enclosure (‘spatial reorientation’)^17–19^. Remembering goal locations relative to boundaries recruits the hippocampus^20^, and humans remember locations close to boundaries better than those far from them^21, 22^.

Within the hippocampal network, an animal’s location relative to boundaries is signalled by different classes of spatially-tuned neuron. Border cells, in the medial entorhinal cortex (mEC), fire when an animal is in close proximity to a boundary, usually in one specific allocentric direction^23, 24^. In the subiculum, a subset of neurons shows spatial firing consistent with that described by the boundary-vector cell (BVC) model^25^, which predicted the existence of BVC neurons, i.e., neurons signalling allocentric distance and direction to boundaries^10, 26^. Recent studies suggest that signalling an organism’s position relative to boundaries may be a common neural mechanism for cognitive mapping, across vertebrates^21, 27, 28^.

Environment boundaries serve as a foundational input to the cognitive map of space during post-natal development: before grid cells emerge (before post-natal day 21), place cells are more stable and accurate when an animal occupies locations close to a boundary^29^.

Border Cells, which may act as a source of stable spatial input to developing place cells, are present in the mEC from P17 onwards^30^. However, how neural responses to boundaries develop in the subiculum remains unknown. Mapping the development of the subiculum is key to understanding hippocampal development overall: the subiculum encodes location, speed, direction, axis of travel and task relevant variables^31–33^, as well as position with respect to boundaries and objects^26, 34^. This information is distributed to multiple brain regions including retrosplenial cortex, nucleus accumbens and anterior thalamic nuclei^35, 36^. Patterns of molecular development indicate that hippocampal maturation recapitulates information flow along the tri-synaptic loop^37^, predicting late maturation of the subiculum. However, to date, developing subiculum neural responses have not been studied in behaving animals.

The goals of this study were to provide the first characterisation of subiculum BVC properties during post-natal development, to understand the nature of the subiculum representation of boundaries, and how this changes through an animal’s lifespan. We categorise subiculum neurons as BVCs by fitting idealised BVC firing rate maps^25^ to neural firing rate maps, obtained by recording the activity of subiculum neurons as rats explored a familiar, square-walled, open field environment. Manipulations of environment geometry revealed that the firing properties of adult BVCs departed from their canonical definition in two ways. Firstly, directional tunings are uniformly distributed in a circle, but cluster in alignment with wall orientations in a square environment, demonstrating an influence of environment geometry on BVC receptive fields. Secondly, insertion of barriers into the open field produces not only a replication of the principal field, but also an inhibition of firing on the opposite side of the barrier, indicating a role for boundary-driven inhibition in reorganising BVC firing fields.

As predicted on the basis of previous observations^37^, the development of precise and stable BVC firing, as well as adult-like responses to inserted boundaries, are slower than those observed in upstream hippocampal areas. However, the fundamental geometry of the distribution of BVC receptive fields, including the influence of geometry on directional tunings in square environments, is observable at the earliest ages tested, suggesting that these are inherent features of the hippocampal coding of space.

## Results

We recorded 575 neurons from the subiculum of 4 adult rats, and 1183 neurons from the subiculum of 17 developing (P16-P25) rats, as they explored a square, walled, open field environment (for tetrode positions see Supplemental Figure 1). To identify BVCs in the open field, we used an exhaustive search for the best fit for each neuronal firing rate map from a set of idealised BVC firing rate maps, constructed following the model described in^25^. The model parameters which were varied to create the test set of idealised BVCs were: the directional tuning, Φ; the distance tuning, *d*; and the width of the distance tuning curve, σ_0_ (Figure 1A, see Methods). If the correlation between the firing rate map and the best fitting idealised BVC was greater than a threshold derived from fitting spatially-shuffled data, the neuron was identified as a BVC (Figure 1B). Additionally, BVCs were required to convey spatial information greater than a threshold derived from spatially-shuffled data (following ^30^; see methods). Figure 1C shows representative examples of classified BVCs in adults and developing rats, with a range of goodness-of-fit from high (top) to just crossing classification threshold (bottom) illustrated for each age group.

**Figure 1.**
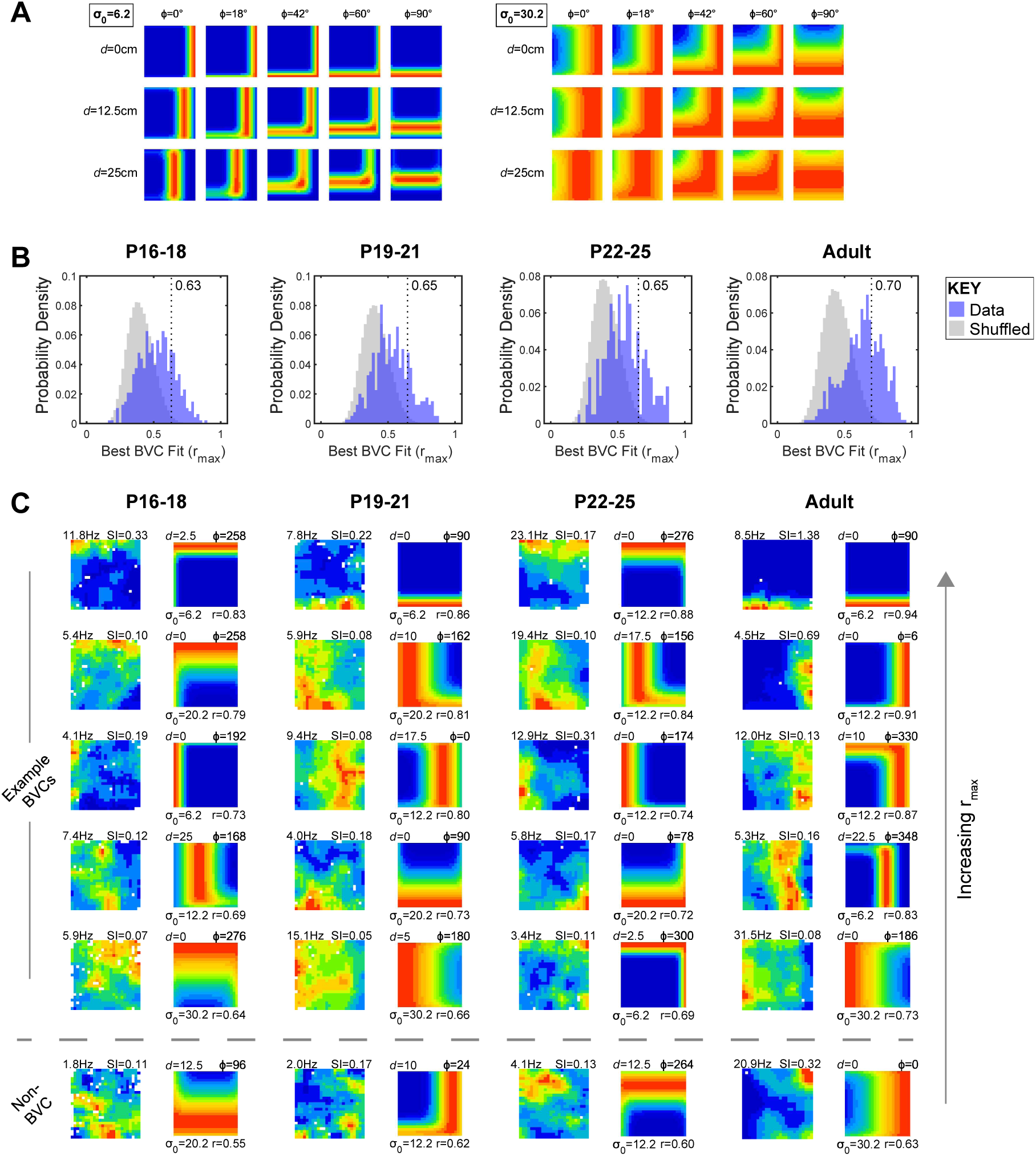
Classification of subiculum neurons as BVCs. **(A)** Example idealised BVC firing rate maps, for different receptive field tunings. Left and right panels show BVCs smallest and largest values of σ_0_ (respectively); within each panel, rows show BVCs differing in *d*, columns show BVCs differing in φ. **(B)** Distributions of correlations between neuronal firing rate maps and the best-fitting BVC map (r_max_) at different ages. Blue histograms show subiculum data r_max_, grey histograms show r_max_ based on shuffled data. Vertical dashed lines show the threshold r_max_ for BVC classification, defined as the 99^th^ percentile of the spike-shifted r_max_ distributions, within each age group. **(C)** Neuronal firing rate maps and respective best-fit BVC model maps, for five example BVCs and one example non-BVC, from each age group. Each row shows example BVCs, each column of paired maps is comprised of the neuronal firing rate map (left) and the best-fit BVC model map (right). For both maps, hot colours indicate high firing rates. Columns of paired maps show data from different ages (cells differ across age groups). Text adjacent to the neuronal firing rate map shows peak rate (top left, Hz) and Spatial Information (‘SI’; top right). Text adjacent to model map shows the *d* (top left), φ (top right), σ_0_ (bottom left) tuning parameters of the BVC, and r_max_ (bottom right). For each age group, representative examples are shown from differing levels of r_max_, sorted from highest (top) to lowest above classification threshold (bottom). The bottom row shows an example cell from each age group which fell below the threshold for BVC classification.

The number of neurons defined as BVCs was significantly greater than expected by chance, at all ages (Figure 2A; Binomial test: p<0.001, all groups). The proportion of BVCs identified in each post-natal group was significantly lower than that obtained in adults (Z-test Adult vs P22-25; Z=3.7, p<0.001), though there was no change in the prevalence of BVCs across the post-natal age range studied (Z-test P16-18 vs P22-25; Z=1.53, p=0.13). Firing rate maps (Fig 1C) show that developing BVCs appear less spatially specific, with greater deviation from their best-fit idealised BVC. Furthermore, comparison across two consecutive trials shows decreased spatial stability in developing BVCs (Figure 2B). The development of the spatial specificity and stability of BVCs was quantified using spatial information, BVC fit r_max_ (see Methods), intra- and inter-trial stability (Fig 2C-F), respectively. All scores increased with age (1-way ANOVA age: Spatial Information, F(_3,415_)=17, p<0.001; BVC r_max_, F(_3,415_)=57, p<0.001; Inter-trial stability, F(_3,415_)=84, p<0.001; Intra-trial stability, F(_3,415_)=15, p<0.001). With the exception of Spatial Information, all measures increased across the pre- and post-weanling period (Tukey HSD P16-18 vs P22-25: Spatial Information p=0.72, all other measures p<0.001). Furthermore, all four measures remained significantly lower in developing animals, including post-weanlings, than in adult rats (Tukey HSD P22-25 vs Adult: all measures p<0.001). In summary, although subiculum BVCs are present in pre-weanling animals, adult- like stability and spatial specificity emerge late in development (>P25).

**Figure 2.**
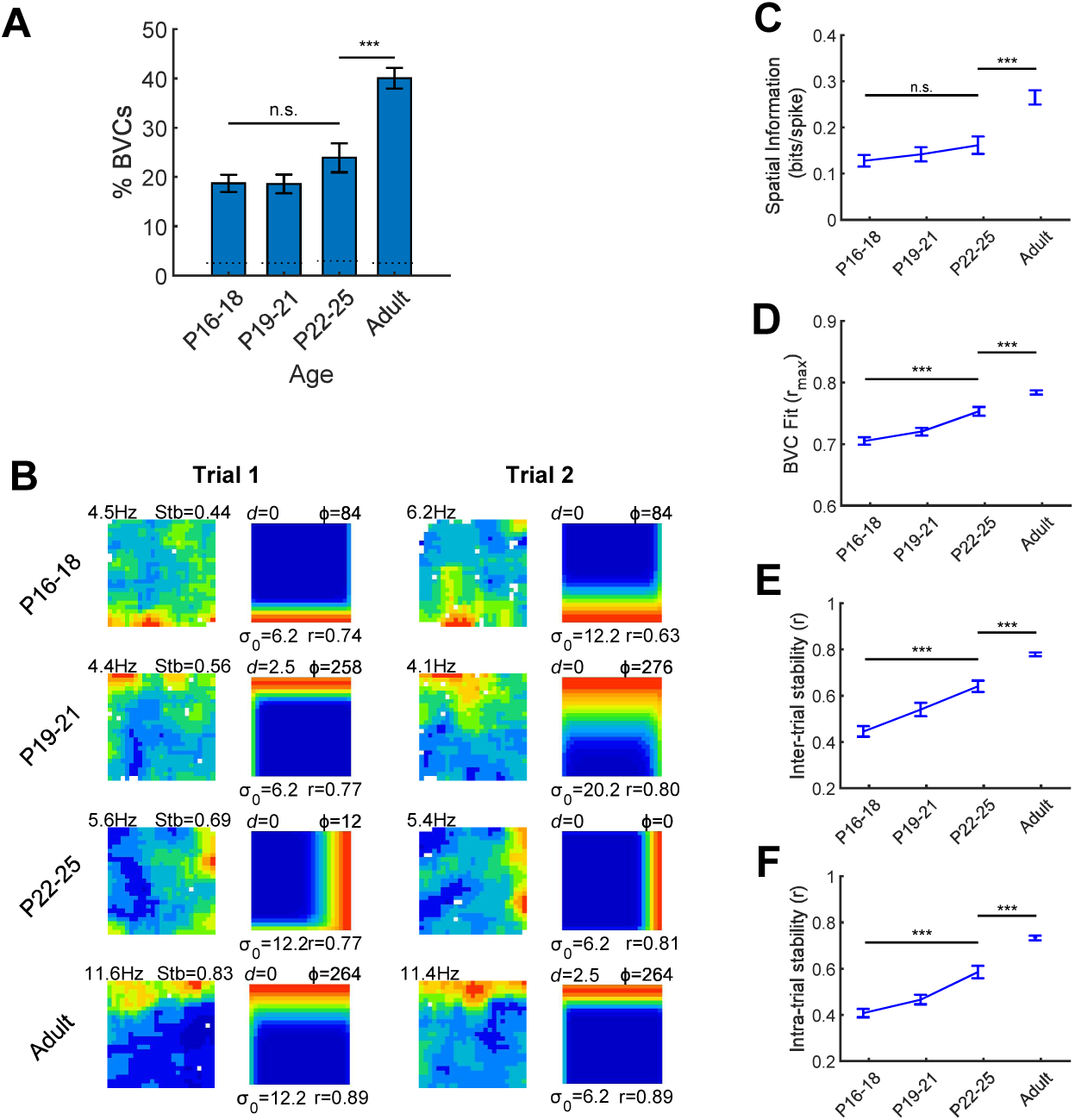
Prevalence, spatial specificity and spatial stability of BVC firing, across development. **(A)** Prevalence of BVCs (percentage of all subiculum neurons classified as BVCs in each age group). Error bars show 95%confidence interval of the proportion. Horizontal dashed lines show 95% confidence level for the number of BVCs exceeding that expected by chance. *** indicates significant differences at p<0.001 level. **(B)** Example BVCs recorded across two consecutive trials, showing increases in inter-trial stability with age. Each row shows a BVC, each column of paired maps shows the neuronal rate map (left) and best-fit model map (right) for one recording trial. Text adjacent to neuronal firing rate map shows peak firing rate (top-left) and inter-trial stability (top right). Model map format as for Figure 1C. Inter-trial stabilities for each example lie within SEM of mean, for respective age group. (**C-F**) Mean (±SEM) values for BVC r_max_ (C), Spatial Information (D), inter-trial stability (E) and intra-trial stability (F), for all BVCs in each age group. *** indicates difference significant at p<0.001 level.

We examined the distributions of spatial tuning parameters (*d*, σ_0_ and Φ) for BVCs recorded at each age. At all ages, distance tunings (*d*) were highly skewed towards short distances, although tunings up to half of the arena width were observed (Figure 3A). The median *d* of recorded BVCs did not change significantly with age (Figure 3B; Kruskal-Wallis test, Age: χ^2^ =5.54, p=0.13). The tuning field widths of BVCs (σ_0_) were distributed over all four tested levels, at all ages (Figure 3C), and the median σ_0_ did not change significantly with age (Figure 3D; Kruskal-Wallis test Age: χ^2^ =5.51, p=0.14). The distribution of directional tunings (Φ) exhibited a striking departure from a uniform distribution: Φ values were strongly clustered at the cardinal points of the compass, aligned to the orientations of the arena walls, in all age groups (Figure 3E). We quantified the four-fold symmetrical clustering using the Rayleigh test on quadrupled, wrapped Φ values (see methods). At all ages, Φ showed a significant four-fold departure from uniformity (P16-18, z=13.4, p<0.001; P19-21, z=16.7, p<0.001; P22-25, z=9.9, p<0.001; Adult, z=57.4, p<0.001). To further compare four-fold clustering across ages, we quantified the proportion of BVCs with Φ oriented ±12° of a cardinal compass point. As expected, this proportion was well above chance, and did not significantly change during development (Figure 3F; χ^2^=1.87, p=0.60).

**Figure 3.**
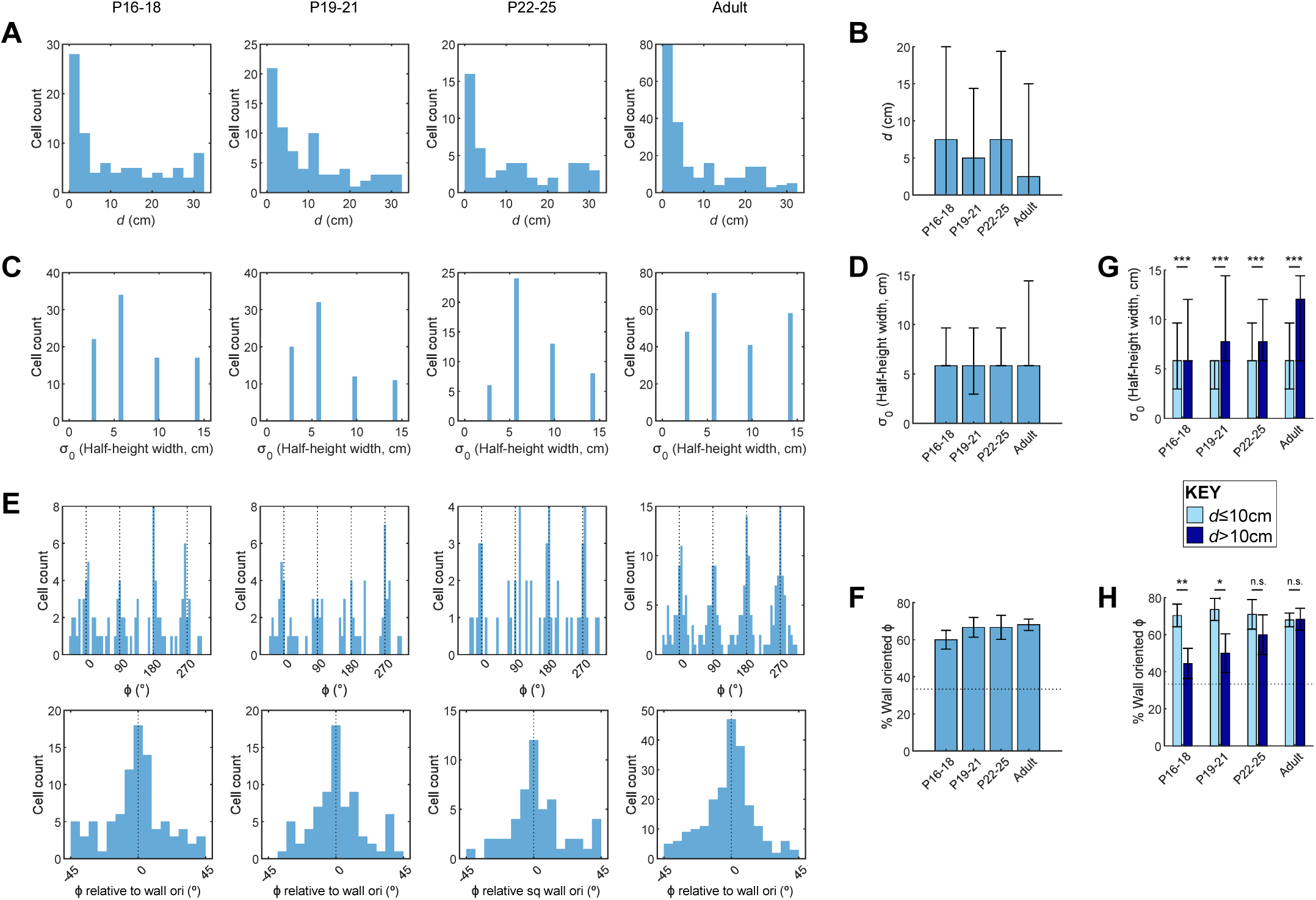
Characterization of BVC tunings for all subiculum BVCs, in each age group. **(A)** Histograms showing all *d* tunings in each age group. **(B)** Median *d* tunings for each age group. Error bars show 25^th^ and 75^th^ percentiles of *d*. **(C)** Counts of σ_0_ tunings in each age group. σ_0_ tunings are expressed as the half-height width of a BVC field with given σ_0_, assuming *d*=0 and φ=0. **(D)** Median σ_0_ tunings for each age group. Error bars show 25^th^ and 75^th^ percentiles of σ_0_. **(E)** Top row: Histograms showing all φ tunings in age group. Black dashed lines show values of φ oriented towards arena walls. Bottom row: histograms of all φ tunings mapped onto one 90° quadrant. 0° indicates φ tunings aligned to a wall. **(F)** Proportion of BVCs with φ oriented towards walls (±12°), in each age group. Error bars show 95% confidence interval. Horizontal dashed line shows expected proportion, assuming a circularly uniform distribution of φ. **(G)** Median σ_0_ tunings for each age, split according to short- and long-range *d* tunings (≤10cm, >10cm, respectively). Error bars show 25^th^ and 75^th^ percentiles of σ_0_. **(H)** Proportion of BVCs with φ oriented towards walls (±12°), for each age group and split according to short- and long-range *d* tunings. Error bars show 95% confidence interval for the proportion. *indicates significant differences at p=0.05 level, ** at p=0.01 level.

Following this, we tested whether the tuning characteristics of BVCs change depending on their preferred firing distance from a boundary, by splitting BVCs into short- and long-range distance tuning groups (*d*≤10cm and *d*>10cm, respectively). Median σ_0_ levels were significantly greater for long-range than short-range BVCs, at all ages (Figure 3G; Wilcoxon test long vs short, p<0.001 for all age groups). The clustering of Φ at wall orientations did not differ between long- and short-range BVCs in adults, but did in developing animals, with Φ being significantly more clustered at wall orientations for short-range BVCs, up until P21 (Figure 3H; χ^2^ Age*Range: χ^2^=20.4, p=0.025; Z-test for proportions short vs long: P16-18, p=0.014; P19-21, p=0.049; P22-25, p=0.42; Adult, p=0.96). Overall, therefore, the geometry of the population of BVC receptive fields is unchanged between early development and adulthood, with the exception that the striking influence of environment walls on directional tunings does not extend to long-range BVCs, until after weaning.

To test whether the clustering of Φ tunings in the square is caused by the geometry of environment boundaries, we exposed the rats to a circular open arena (diameter 80cm). The distributions of Φ and *d* tunings in both square and circle are illustrated in Figure 4A, which shows a clear contrast between the distribution of Φ tunings in the square (clustered at cardinal compass points) and those in the circle, which possess an apparently uniform angular distribution. Indeed, BVC directional tunings in the circular arena showed no significant 4-fold radial symmetry or unimodal departure from uniformity (Figure 4B; Rayleigh test quadrupled Φ: P16-18, z=0.6, p=0.56; P19-21, z=2.7, p=0.07; P22-25, z=0.8, p=0.47; Adult, z=1.8, p=0.17; Rayleigh test unimodal: P16-18, z=2.7, p=0.067; P19-21, z=2.7, p=0.060; P22-25, z=0.9, p=0.40; Adult, z=0.0, p=0.95). Inspection of the rate maps reveals that Φ tunings sometimes rotated between square and circle in the laboratory reference frame, likely due to the lack of common extra-maze cues across these environments (Figure 4C; see Methods). Within each ensemble, rotations of individual BVC Φ tunings were clustered around the ensemble mean rotation for both pups and adults (Figure 4D; Concentration parameter kappa: P16-25, k=1.38; Adult, k=1.87; K-test for differing Kappa [Pup vs Adult]: f=1.37, p=0.19), indicating that BVC ensemble rotations were approximately coherent. However, relative Φ rotations were not fully rigid across co-recorded cells, with Φ offsets between cell pairs often changing within the range ±45° (see examples Figure 4C; median absolute rotation relative to mean: P16-25, 30°; Adult, 22°). This partial plasticity in relative Φ offsets is consistent with a reorganisation of ensemble receptive fields between circle and square, producing the 4-fold clustering observed in the latter environment.

**Figure 4.**
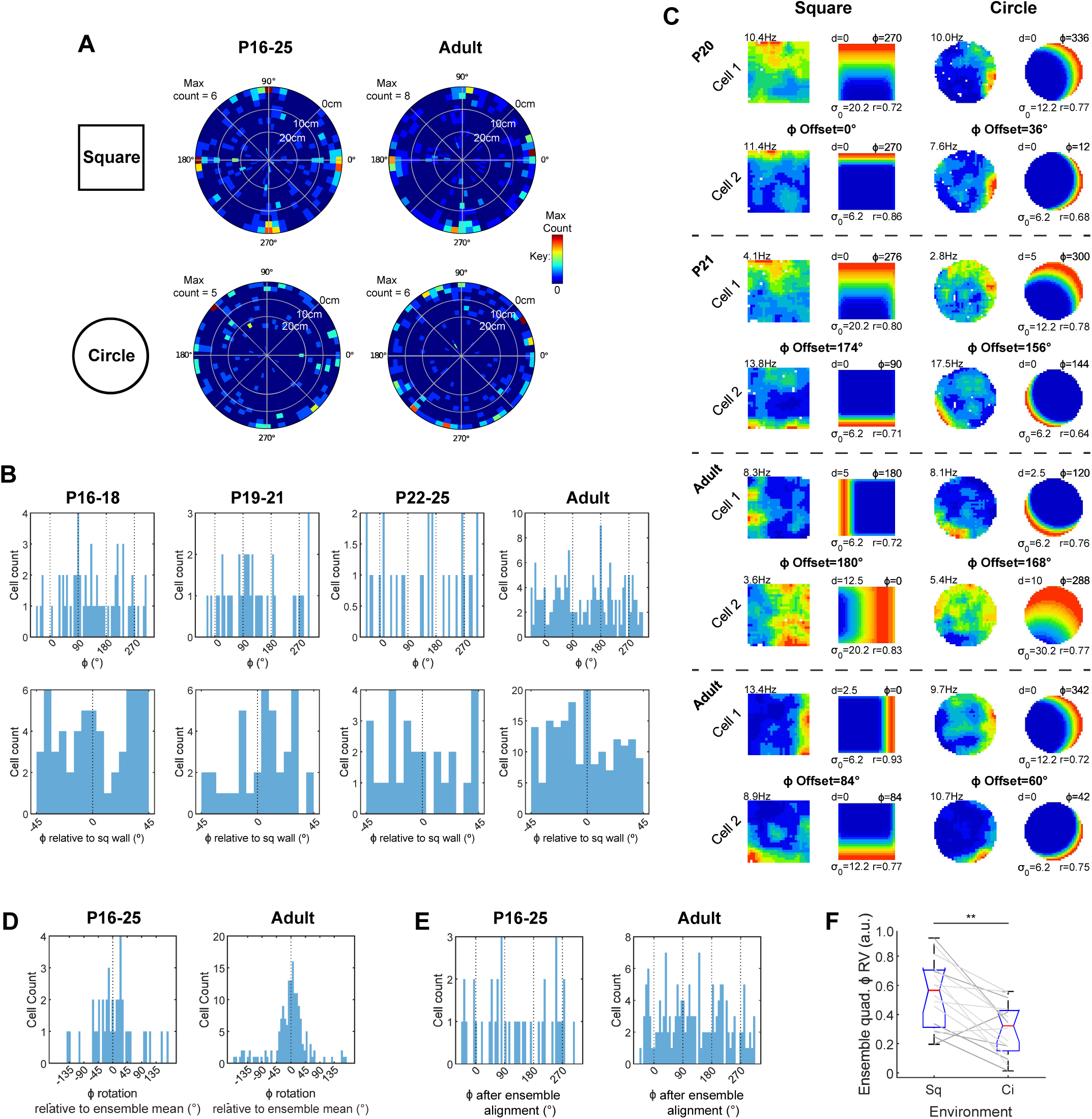
Characterisation of BVC φ tunings in a circular arena. **(A)** Two-dimensional (polar) histograms showing distributions of BVC *d* and φ in the square and circular arenas, in adults and developing rats. Bins colours show histogram counts (see Key), angular axis shows φ, radial axis shows *d.* Radial axis is reversed such that low *d* is outermost, to enhance visual clarity of clustering of φ tunings in the square arena. **(B)** Top row: Histograms showing all φ tunings in age group, in the circular arena. Black dashed lines show values of φ orientated towards the square arena walls. Bottom row: histograms of all φ tunings mapped onto one 90° quadrant. 0° indicates φ tunings aligned to the square environment walls, in the laboratory reference frame. **(C)** Neuronal firing rate maps and respective best-fit BVC model maps for four pairs of simultaneously recorded BVCs, in the square and circular arenas. Each row shows one BVC, dashed lines delineate simultaneously recorded pairs. Text top left of neuronal map shows peak firing rate. Model map format as for Figure 1C. Text between rows shows offsets of φ tuning between co-recorded pairs in square and circle. **(D)** Histograms showing rotations of BVC φ tunings, relative to the mean φ rotation for their ensemble (ensembles of ≥5 BVCs only). Left panel shows all developing rats, right panel shows adults. **(E)** Histograms showing distribution of φ tunings after alignment to common directional reference frame, by subtraction of ensemble mean φ rotation from individual BVC φ rotations (ensembles of ≥5 BVCs only). **(F)** Rayleigh vector lengths derived from quadrupled wrapped φ (quad-φ RV), for each ensemble with ≥5 BVCs. Box plots show distribution of quad-φ RV in square and circle arenas, grey lines show change in quad-φ RV for each ensemble, between square and circle.

The rotation of BVC ensembles between square and circle raises an alternative explanation for our results: that 90° Φ clustering is present in the circle within each ensemble, but the orientation of these clusters is inconsistently aligned across ensembles, obscuring the clustering at the population level. To test this possibility, we corrected the Φ tuning for each BVC by subtracting the mean ensemble rotation (for ensembles with ≥5 co-recorded BVCs). Even following this correction procedure, the distribution of Φ tunings, in circular environments, showed no significant 4-fold departure from uniformity (Figure 4E, Rayleigh test quadrupled Φ: P16-25, z=2.0, p=0.14; Adult, z=2.0, p=0.61). Furthermore, even within each co-recorded ensemble (≥5 co-recorded BVCs), BVC Φ tuning distributions showed a significant reduction of 4-fold symmetry when moving from square to circle, as shown by the length of the Rayleigh vector derived from wrapped, quadrupled Φ (Figure 4F; Wilcoxon Test; p=0.002). Features of boundary geometry present specifically in the square, as opposed to the circle, are therefore most likely responsible for the observed 90° clustered distribution of BVC Φ tunings in square environments.

An important characteristic of boundary-driven firing is that the introduction of a new boundary, such as a barrier, into the environment causes a new firing field to emerge^9,23^. To assess the development of barrier-driven firing, a barrier (oriented EW or NS) was inserted into the square arena, and we quantified whether BVC firing increased on the distal side (relative to the existing BVC field), as compared to the proximal side of the barrier (following^30^; see Methods). At all ages, barrier insertion caused an increase in absolute firing rate on the distal side (Figure 5), though the difference between distal and proximal firing rates did not reach significance until P19 (Mixed ANOVA: Age*Barr Side F_(3,202)_=17.1, p<0.001; SME Dist vs Prox: P16-18, p=0.057, P19-21, p=0.014, P22-25, p=0.004, Adult p<0.001). However, when changes in firing rate were normalised to overall firing rates in the barrier and baseline trials, the difference between proximal and distal side firing was significant at all ages (Figure 5C; Mixed ANOVA: Age*Barr Side F_(3,202)_=27.2, p<0.001; SME Dist vs Prox: p<0.001 all age groups), suggesting that part of the slow development of barrier-driven fields is due to low overall firing rates. Despite this, it remains the case that barrier-driven responses were weaker in all juvenile groups than in adults, both in terms of absolute and normalised firing (SME Dist_(P22-25)_ vs Dist_(Adult)_: Absolute rate, p=0.017; Normalised rate, p=0.015).

**Figure 5.**
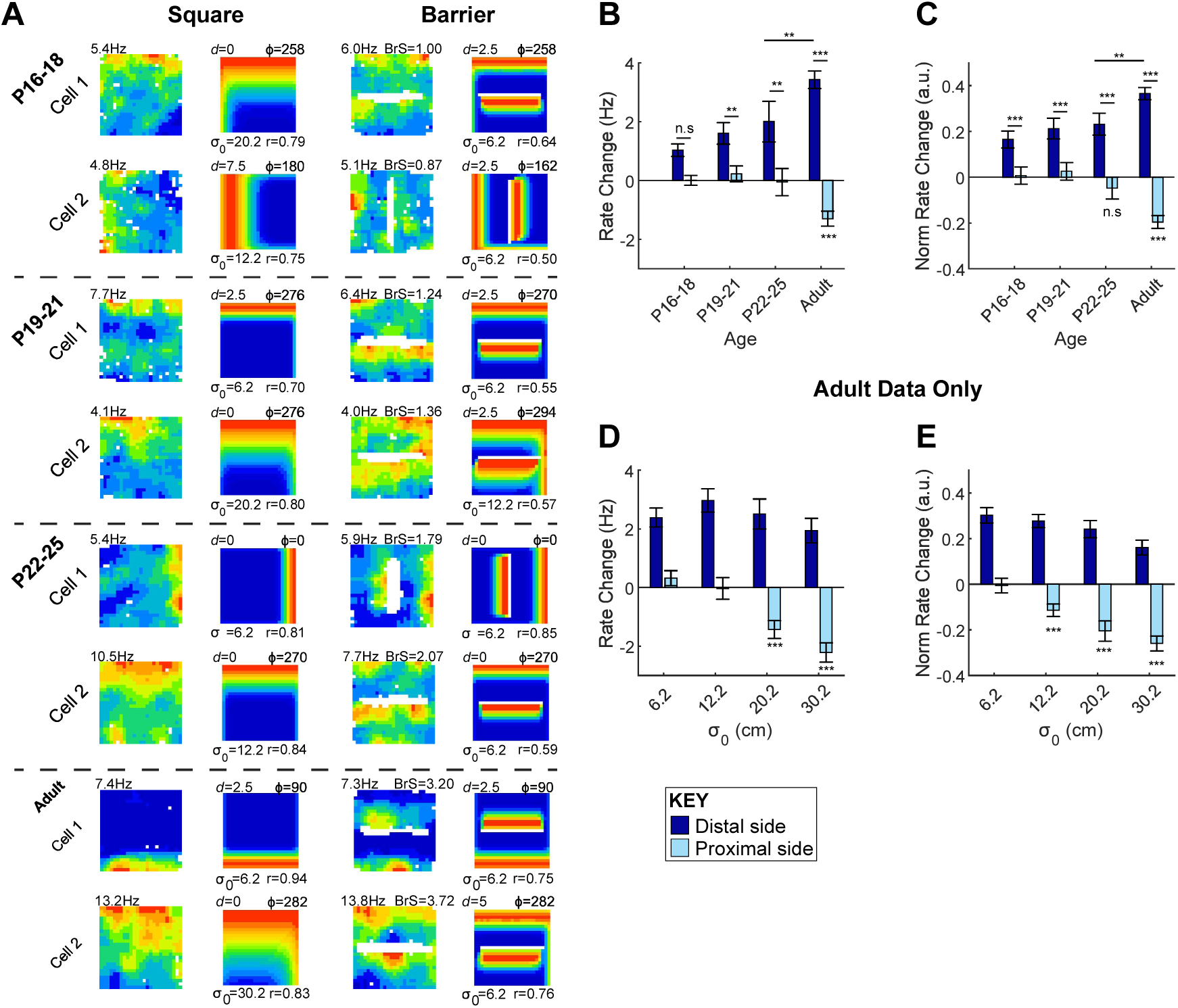
Development of BVC response to inserted boundaries. **(A)** Neuronal firing rate maps and best-fit BVC model maps for eight example BVCs, in the square arena (left columns) and following the insertion of a barrier (right columns). Each row shows one BVC, dashed lines delineate cells recorded at different ages. Text adjacent to neuronal firing rate map shows peak rate (top-left, Hz) or the barrier rate change score (top-right, ‘BrS’). Model map format as for Figure 1C. **(B)** Mean barrier rate change score (±SEM) for BVCs in each age group. Dark blue bars show rate changes on distal side of the barrier, light blue bars on the proximal side of the barrier. **(C)** Mean barrier rate change score (±SEM) for BVCs in each age group, normalised to overall rate across square and barrier trials. **(D)** Mean barrier rate change score (±SEM) for adult BVCs, separated by σ_0_ of best-fit BVC in baseline trial. **(E)** Mean normalised barrier rate change score (±SEM) for adult BVCs, separated by σ_0_ of best-fit BVC in baseline trial. *indicates significant differences at p=0.05 level, ** at p=0.01 level, *** at p=0.001 level. Asterisks directly under light blue bars indicate significance of difference from rate change=0.

Notably, in adults, barrier insertion causes a significant inhibition of firing rate on the proximal side of the barrier, relative to baseline level, indicating that barrier insertion causes a reorganisation of the BVC field that goes beyond simply replicating the principal firing field (1- sample T-Test versus no change: Absolute rate, p<0.001, Normalised rate, p=0.001). However, this reduction in firing is not observed in any development group (p≤0.34, all groups). In adults, proximal-side inhibition was stronger in BVCs with broader-tuned receptive fields (Figure 5A c.f. bottom two rows, 5D-E; 1-way ANOVA Best fit model σ_0_: Absolute Proximal Rate, F_(3,279)_=12.8, p<0.001; Normalised Proximal Rate, F_(3,279)_=11.8, p<0.001) and was only consistently significant (across both absolute and normalised rate measures) for cells falling in the two broadest σ_0_ categories (1-sample T-Test versus no change: Absolute rate, σ_0_=6.2, p=0.90; σ_0_=12.2, p=0.462; σ_0_=20.2, p<0.001; σ_0_=30.2, p<0.001; Normalised rate, σ_0_=6.2, p=0.43; σ_0_=12.2, p<0.001; σ_0_=20.2, p<0.001; σ_0_=30.2, p<0.001). Where an adult BVC has a broader baseline field, this is consistently inhibited by barrier insertion, on the proximal barrier side. Splitting developing BVCs by best-fit σ_0_ revealed a trend for broader fields to be inhibited, exclusively in post-weanling pups, but this effect did not reach statistical significance (see figure S2). In summary, while excitatory responses to inserted barriers are observed in subiculum BVCs from P16, the inhibitory component of the BVC response observed in fully mature BVCs does not emerge until much later in development.

We have shown that subiculum BVCs, although present at P16, continue to mature late into development (>P16), with spatial tuning, stability, and barrier responses all remaining immature as late as P25. This is in contrast to previous reports of mEC border cells^30^ which showed no change in either spatial tuning or stability between P17 and adulthood. These contrasting results could be due to the faster maturation of mEC (as suggested by^37^). However, an alternative possibility is that the boundary-responsive cells captured by the BVC measure mature more slowly than those defined using the border score (which is biased towards selecting fields near to boundaries). To rule out this latter possibility, we re-analysed the data described by^30^, and quantified the development of boundary-driven firing using either the border score, or the BVC classification method (as described here) to define boundary-responsive neurons.

Figure 6A shows the proportion of mEC neurons classified as either a border cell, a BVC or both. Distributions of shuffled and actual BVC fits to the data are shown in figure S3A. As previously reported, there is no difference between the proportions of mEC border cells in adults and developing rats (Z-test proportions adult vs developing: Z=0.33, p=0.74), and the same is true for the proportion of BVCs (Z=-1.02, p=0.31). At all ages, there is considerable overlap between those boundary-responsive neurons classified as border cells and BVCs, but the proportion of these “intersect” cells does not change between adults and developing rats (Z=-1.1, p=0.27). Figure 6B shows examples of neurons which were classified as BVCs, Border Cells, or as both types simultaneously. Cells classified as both BVCs and border cells show clear firing fields close to, and extending along, boundaries. Cells selected uniquely by the BVC measure show broader firing fields with firing extending away from walls. Neurons classified as border cells, but not BVCs, by contrast, exhibit multiple firing fields which are close to walls, but do not cover their full extent. For all cell types, the qualitatively-judged firing properties do not change with age.

**Figure 6.**
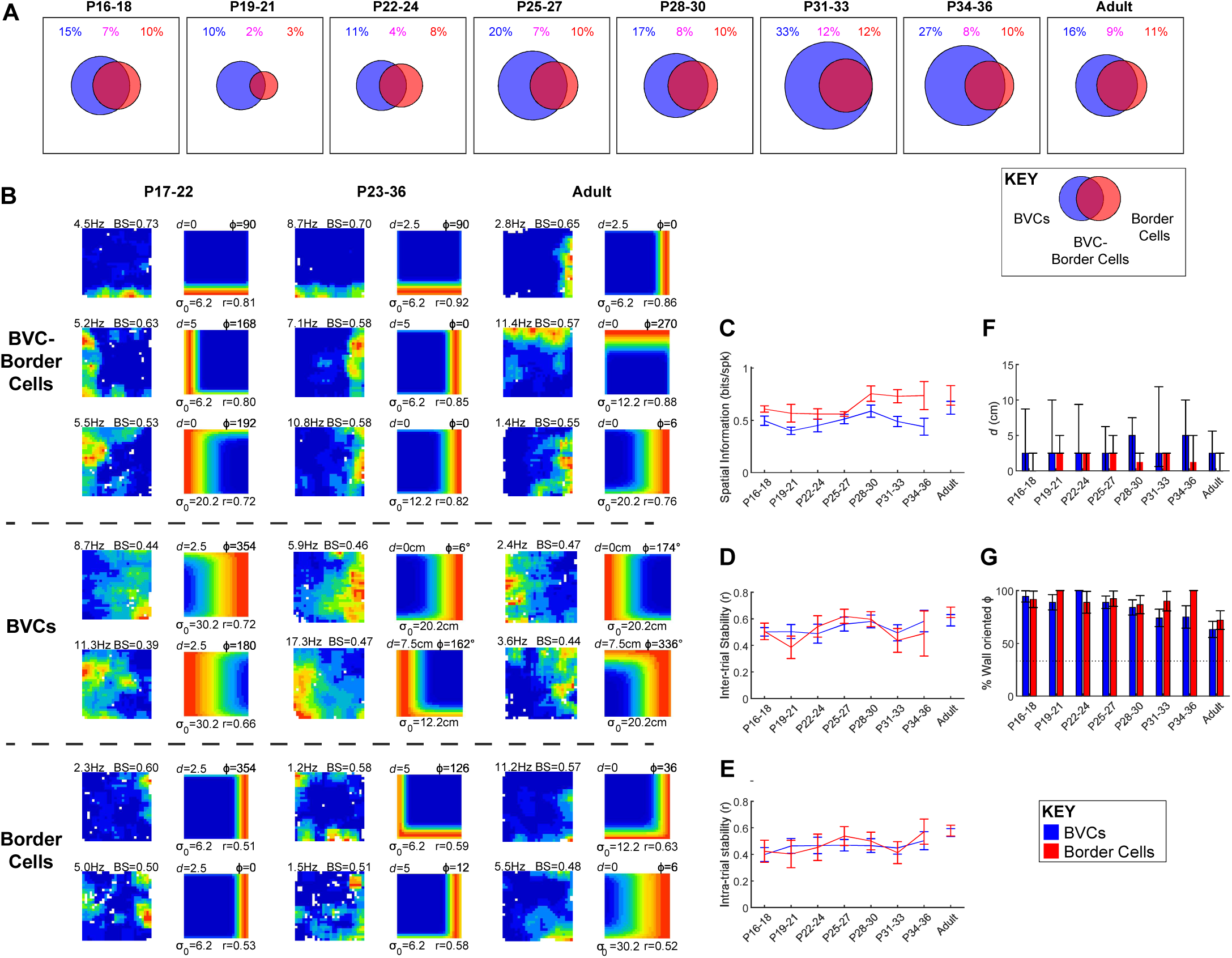
Comparison of BVCs and border cells recorded in medial Entorhinal Cortex. **(A)** Venn diagrams showing the proportion of mEC cells classified as BVCs (blue), Border Cells (red), or both cell types (overlap region), for each age group. Coloured text above circles shows corresponding numerical percentages. Circles are scaled such that the square bounding box represents 100% of cells recorded in age group. **(B)** Neuronal firing rate maps and best-fitting BVC model maps, for example BVCs and Border Cells. Each row shows example cell, columns of paired maps show data from different age groups (cells differ across age groups). Text adjacent to the neuronal firing rate map shows peak rate (top left, Hz) and Border Score (‘BS’; top right). Model map format as for figure 1C. Top three rows show cells classified as both BVCs and Border Cells, middle two rows cells classified as BVCs but not Border Cells, bottom two rows cells classified as Border Cells but not BVCs. (**C-E**) Mean (±SEM) values of Spatial Information (C), inter-trial stability (D) and intra-trial stability (E), for BVCs (blue line) and Border cells (red line) in each age group. Cells classified as both BVCs and Border Cells are included in both groups. **(F)** Median *d* tunings for BVCs (blue bars) and Border Cells (red bars), in each age group. Error bars show 25^th^ and 75^th^ percentiles of *d* for each group. **(G)** Proportion of BVCs (blue bars) and Border Cells (red bars) with φ oriented towards walls (±12°), in each age group. Error bars show 95% confidence interval for the proportion. Horizontal dashed line shows proportion expected, assuming a circularly uniform distribution of φ

We confirmed the precocious maturity of mEC boundary-related firing by quantifying the spatial information and stability of the classified cells. The spatial tuning and spatial stability of mEC BVCs does not change during development (Figure 6C-E, ANOVA Age: Spatial Information, F_(7,166)_=1.75, p=0.10; Inter-trial stability, F_(7,160)_=0.65, p=0.72; Intra-trial stability, F_(7,166)_=1.17, p=0.33), and furthermore, we confirmed that neither does the tuning and stability of border cells (Spatial Information, F_(7,84)_=0.05, p=0.40; Inter-trial stability, F_(7,85)_=1.8, p=0.10; Intra-trial stability, F_(7,84)_=0.99, p=0.44). Boundary-responsive firing in the mEC appears, therefore, mature from P17, irrespective of the measure used to quantify it. Similarly, the tuning properties of mEC BVCs do not change throughout development: though BVCs have longer distance tunings than border cells (Figure 6F, Figure S3B, S3E; Wilcoxon test BVC vs Border Cell all ages: p<0.001), the median distance tunings of either does not change during development (Kruskal-Wallis Age: BVCs, χ^2^ =5.9, p=0.55; Border Cells, χ^2^ =11.8, p=0.11). The proportion of angular tunings oriented to walls is significantly higher than chance, at all ages (Figure 6G, Figure S3D, S3G), and does not change with age (χ^2^ Age: BVCs, χ^2^=40, p=0.25; Border Cells, χ^2^=48, p=0.24). The distortion of boundary-responsive angular symmetry is also apparent in mEC boundary responsive neurons, from early in development (though responses to a circular arena were not tested within this dataset).

## Discussion

This study is the first to define a population of BVCs in the subiculum and characterise their spatial tuning. Previous reports of BVCs in the subiculum did not use a formal objective procedure to define a BVC^10, 26^, and the method used to define Border Cells in mEC^23, 30^ selectively weights firing fields in close proximity to boundaries, and is not well-suited to capture the broader and vectorial nature of spatial firing in the subiculum. The BVC model fit method used here captures both long- and short-range boundary tunings, and our parallel analysis of mEC and subiculum data confirms that mEC boundary-driven cells respond selectively closer to walls, whereas in the subiculum a broader range of distance tunings are present (cf Figure 3B, 6F). Subiculum neurons whose spatial tuning closely fits that of idealised BVCs have previously been reported^33^, using a similar method to our own, though this study did not provide any further characterisation of the BVCs detected. The goodness-of-fit r-values appeared markedly lower in that study than those reported here: this may be explained by the fact that^33^ used a wall-less arena, which can reduce the spatial specificity of BVCs^9^.

The model-fitting method used here allowed us to characterise the population of detected BVCs in terms of the preferred distance-to-boundary (*d*), allocentric angle-to-boundary (Φ) and receptive field width (σ_0_) of the idealised BVCs fitted to neural data. Here we show that preferred distance tunings are biased towards short-range tunings (modal *d* was 0 cm, at all ages), but the full range of possible preferred distances were detected, up to the arena radius. The preferred distance-to-boundary tunings described here are smaller than recently reported object-responsive subiculum neurons^34^ (Vector Trace Cells 14.1±1.0cm [Mean±SEM], Non-trace cells 11.2±0.4cm; Adult BVCs in present study, 7.5±0.6cm), which may reflect the smaller arena used in this study (62.5cm, versus 100cm) preventing the detection of longer-range BVCs. Nonetheless, this study confirms the vectorial nature of boundary coding in the subiculum and builds on previous reports of object vector coding^34, 38–40^ to suggest that the vectorial coding of allocentric space is fundamental to hippocampal spatial cognition.

A striking feature of BVC tunings is the non-uniformity of the receptive field preferred angles in a square arena, which were strongly clustered with a four-fold circular symmetry, aligned with walls of the square environment (as predicted by^41^), whereas this four-fold clustering is not present in a circular environment. BVC receptive fields are unlikely, therefore, to solely represent an allocentric vector to boundaries^25^, fixed across all environments, but are instead further modulated by the geometric features of those boundaries (e.g. straight walls and corners). This result is consistent with both the well-established finding that the geometry of walls and corners influences spatial memory^17–19, 22^, and that corners in particular may be an important class of geometric cue, with acute corners being more salient than obtuse corners^42, 43^. The modulation of BVC firing by boundary geometry, and corners in particular, may therefore represent a mechanism by which these geometric features exert effects on behaviour.

The distortion of BVC firing by environment geometry echoes previous findings of distortion of grid cell firing patterns^44, 45^. The early emergence of subiculum BVCs in relation to grid cells^30, 46, 47^ suggests that the influence of geometry on BVCs is independent of its effects on grid cells. Our study confirms that boundary firing in both subiculum and mEC aligns with individual walls, supporting the hypothesis^44^ that grid field shearing may be caused by a combination of grid shrinkage and anchoring of the grid to a sub-set of local environment cues (e.g., one boundary of the environment). A further example of environmental geometry influencing spatially modulated firing is the fact that square environments enforce a four-fold symmetry in the directional tuning curves of egocentric boundary cells in the retrosplenial cortex^48^, which the authors speculate may create boundary-driven distortions in downstream allocentrically-tuned neurons.

Boundary geometry can affect spatial memory in ways that recapitulate its effects on spatially-modulated neurons: distance estimates are distorted in a rhomboid environment, particularly at the narrow end^49^; shrinking or stretching a square environment shifts goal search locations in a way predicted by place field movements^22^. Thus, it is likely that the distortion of BVC fields by geometry is a mechanism by which boundary geometry influences behaviour. The observation that BVC fields cluster along walls in the square environment suggests that the representation of corner locations in the square may be less precise than the equivalent allocentric locations in the circular environment. Consistently with this hypothesis, moving between a square- to circular-bounded watermaze (whilst keeping all extra-maze cues constant) disrupts memory for platform position^50^, whilst shape transfers that preserve local geometry of corners and walls in specific parts of the maze also preserves memory of these locations, in a hippocampally-dependent fashion^51, 52^.

The inhibition of adult BVC firing on the proximal side of an inserted barrier is a further property of BVCs not predicted by the canonical model^25^, which predicted the existence of solely excitatory receptive fields. Inhibitory receptive fields have also previously been described in unpublished work^53^. One explanation for these findings is that BVCs (directly or indirectly) inhibit other BVCs with directionally-opposed receptive fields, analogously to continuous attractor connectivity of head direction cells^54, 55^. This would be consistent with the prediction that BVCs form an attractor representation of allocentric position with respect to boundaries^56^, and also explain the observed rotational coherence of subiculum spatial firing, observed in this and other studies^57^. Indeed, neurons whose firing matches the inverse of a BVC field (including some with interneuron-type waveforms) have previously been reported in the subiculum^9^,

In addition to describing the adult subiculum BVC population, here we also offer the first characterisation of the emergence of subiculum BVCs during post-natal development, and specifically across weaning, when hippocampal learning and memory first emerges^58^. We found that the development of BVCs has both precocious and late-emerging aspects: the spatial stability and specificity of BVCs, and their responses to inserted barriers, develops gradually and slowly, mirroring the previously described gradual emergence of place cell firing in CA1^46, 47, 59^. This pattern of development provides a notable contrast with boundary-driven firing in the mEC, which appears adult-like from P17 onwards^30^ – a finding we have confirmed in our reanalysis of mEC data. These data are consistent with the proposal that maturation of different hippocampal regions proceeds sequentially, along the tri-synaptic loop, with entorhinal areas maturing first, followed by CA3, CA1 and subiculum^37^. The slow maturation of BVC specificity is consistent with the late-emerging ability of boundaries to enable accurate recall of spatial locations^60, 61^, and suggests that BVC development may be a key limiting step in the late emergence of place learning more generally^62, 63^.

In other respects, however, the development of BVCs is precocious. BVC firing is present at the earliest ages tested, and the overall characteristics of BVC spatial tunings, at the population level, appear in adult-like form: average d and σ_0_ do not change with age, and Φ tunings are clustered to align with wall orientations in the square. The core geometrical properties of the BVC representation of space thus appears in an already mature form at the earliest age tested (P17). The only exception to this pattern is the distorting influence of boundaries on BVC directional tunings for BVCs with long distance tunings, which only emerges post-weaning, a finding which echoes increases in place cell stability far from walls at the same age^29^. The presence of a specific geometry of BVC tunings for square environments may be a neural mechanism enabling the recognition of geometric boundary information, thereby underlying the early emergence of spatial reorientation^18, 64, 65^. It is unclear whether this mechanism is innate or experience dependent. We note that the animals in this study were reared in rectangular cages: it remains to be established how BVCs would respond following rearing in circular environments, a manipulation which impairs the use of geometrical information in rodents^64^.

## Acknowledgements

We thank Colin Lever for revision and very helpful discussion of the manuscript. We thank the following funders for supporting this research: Wellcome Trust (Senior Research Fellowship 220886/Z/20/Z to T.W., PhD Studentship to F.RR. and BT), The Royal Society (University Research Fellowships UF150692 and UF100746 to T.W.), Medical Research Council, UK (MR/N026012/1 to TW), European Research Council (Starter Award DEVSPACE and Consolidator Award DEVMEM to F.C.), Biotechnology and Biological Sciences Research Council, UK (grant BB/I021221/1 to F.C.).

## Author Contributions

Conceptualization, T.W. and F.C.; Investigation, L.M., F.R-R.; Formal Analysis & Visualization, L.M., T.W.; Writing – Original Draft, T.W., F.C.; Writing – Review & Editing, T.W., F.C., L.M., F.RR., N.B, C.B., E.I.M., MB.M.; Funding Acquisition, T.W., F.C.; Resources, B.T., C.B., N.B.

(BVC fitting algorithm), T.B., E.I.M., MB.M. (previously published data contributing to figure 6); Supervision, T.W., F.C.

## Methods

### Subjects

Subiculum data was collected from 17 developing male lister hooded rat pups (aged P12-P14 and weighing 24-32g at time of surgery) and 4 male (3-6mo) lister hooded rats. Developing rats were bred in-house and remained with their dams until weaning (P21). Rats were maintained on a 12:12 hour light:dark schedule (with lights off at 10:00). At P4, litters were culled to 8 pups per mother in order to minimise inter-litter variability. Pregnant females were checked at 17:00 daily and if a litter was present, that day was labelled P0. After surgery (see below), each pup was separated from the mother for between 30 minutes and 2 hours each day, to allow for electrophysiological recordings.

### Surgery and electrode implantation

Rats were anaesthetised using 1-2% isoflurane, and 0.15mg/Kg bodyweight buprenorphine. Rats were chronically implanted with microdrives loaded with 8 tetrodes (HM-L coated 90% platinum-10% iridium 17μm diameter wire). Microelectrodes were aimed at the hippocampal subiculum region. In developing rats the co- ordinates used were 4.4 mm posterior and 1.3 mm lateral to Bregma, 2.7 mm below brain surface. For adult rats, the co-ordinates used were 5.4 mm posterior and 1.5 mm lateral to Bregma, 2.7 mm below brain surface. After surgery, rats were placed in a heated chamber until they had fully recovered from the anaesthetic (10 - 30 minutes), and were then returned to the mother and littermates.

### Single-unit recording

Rats were allowed a 1-day postoperative recovery, after which microelectrodes were advanced ventrally by 62-250 µm/day until they reached the subiculum cell layer, identified on the basis of a prominent theta (5-8Hz) LFP rhythm and the presence of theta-modulated pyramidal cell firing, at which point recording sessions began. Single units in the subiculum were defined as excitatory pyramidal cells on the basis of a waveform width >300µs. Single unit data was acquired using an Axona (Herts, UK) DACQ system. LEDs, were used to track the position and directional heading of the animal. Isolation of single units from multi-unit data was performed manually on the basis of peak-to-trough amplitude, using the software package ‘TINT’ (Axona, Herts, UK). Rat position was recorded by tracking two LEDs attached to the headstage amplifier.

### Behavioural testing and environments

Single-unit activity was recorded while rats searched for drops of soya-based infant formula milk randomly scattered on the floor of two different open field arenas: (1) a square-walled (62.5cm sides, 50cm high) light-grey wooden box, placed on a black Perspex floor, (2) a circular-walled (80cm diameter, 50cm high) light-grey wooden box, placed on a black painted wooden floor. From the square arena, distal visual cues were available in the form of the fixed apparatus of the laboratory. The circular arena was placed within a set of closed black curtains, within which there was only one prominent distal visual cue, an A0-sized white card, illuminated by a 20W (incandescent filament) desk lamp. Before recording in the circular arena, rats were carried through the curtains in a closed black box, which was moved directly between the arenas, without being rotated. To test the inter-trial stability of BVCs, two temporally adjacent trials were run in the square open arena. To test responses of subiculum neurons to barriers, a single straight barrier was inserted into the standard square arena, aligned parallel to the arena walls, centred within the open field in both x and y dimensions. In the majority of recording sessions (83% of recorded BVCs), both possible barrier orientations (North-South and East-West) were tested. Otherwise, only one orientation was tested, based on the orientation of BVCs as assessed in the baseline (open field square) trials. Barriers were made of the same material and of the same appearance as the arena walls. The majority of rats were tested using a barrier of dimensions 50cm length, 2.5cm width and 50cm height; 4 developing rats were instead tested using a barrier of 40cm length, 5cm width and 50cm height. Trials were 15 minutes long. Between each trial, the rat rested for a 15 minute inter-trial interval in a 25cm x 25cm holding box containing a heated pad.

### Construction of Firing Rate Maps

All spike and positional data were filtered so as to remove periods of immobility (defined as speed < 2.5cm/s). Following this, positional data were re- scaled to a standardised size and shape, for consistent comparison to BVC model maps. For the square arena, edges of the arena along each wall were defined as the line of camera pixels (each pixel being 2.5mm wide) furthest from the centre of the environment where the summed dwell time was ≥1 sec. Data outside these edges was discarded, and the remaining data was divided into a 25 x 25 grid of evenly spaced spatial bins, each representing an area 2.5cm x 2.5cm. For circular arena data, the centre of the arena was first estimated as being the mid-point between the visited edges of the environment at the cardinal points of the compass, which were defined as the line of camera pixels furthest from the centre where the summed dwell time ≥0.2 sec. Following this, all positional data was rotated by 45° around the estimated centre, and the edges and centre were defined again, following the method described above, for the rotated data. These steps were continued in an iterative fashion until consecutive estimates of the centre converged to within 0.75 cm. Following the definition of the centre of the environment, the edge of the environment was defined as the largest pixel-wide circumference around the centre for which the total summed positional dwell time was ≥1 sec. Data outside these edges was discarded, and the remaining data was divided into a 32 x 32 grid of evenly spaced spatial bins each representing an area 2.5cm x 2.5cm.

Following standardised scaling of the data, total positional dwell time and spike count for the whole trial was calculated for each spatial bin. The binned position dwell time and spike count maps for each cell were then smoothed using a boxcar filter, 5×5 bins.

### Construction and fitting of BVC model maps

Firing rate maps corresponding to the activity of an idealised BVC were defined following ^25^. Briefly, the firing rate *g* of a BVC tuned to respond maximally to the presence of a boundary segment at distance *d* and allocentric direction Φ was defined as:

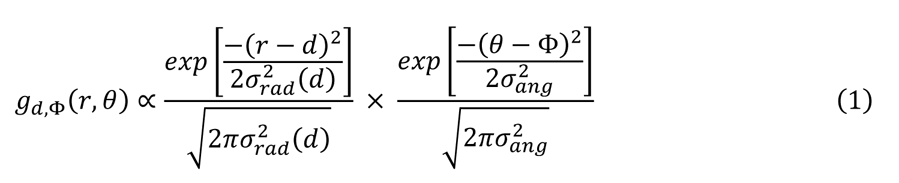

where *r* is the distance and *ɵ* the allocentric direction from the animal to the boundary segment, *a*_*ang*_ is constant, and radial field extent *a*_*rad*_ varies linearly with

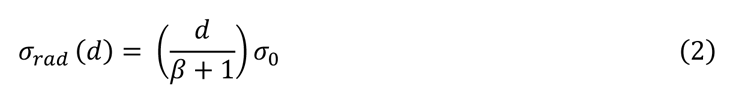

The firing rate *f*(*x, y*), at any position (*x, y*), of a BVC with receptive field *g*_*d,,Φ*_ can therefore be defined by summing equation 1 over all directions *θ* in steps of size *Δθ*:

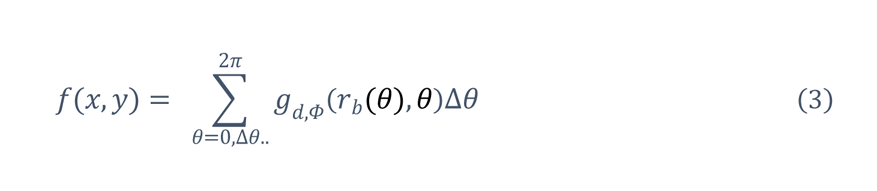

where *r*_*b*_(θ) is the distance to the first boundary segment in direction *r*_*b*_ (if there is no boundary in that direction in that direction is infinite and no firing results). Equation (3) was then evaluated at a series of points corresponding to the centres of an evenly spaced 25×25 grid of 2.5cm x 2.5cm spatial bins to give the firing rate map for the BVC. The boundary of the environment was defined as falling at the edge of the outermost bins of the grid. For the circular arena, the boundary was defined as a 80cm wide circle, centred within a 32×32 grid of 2.5cm × 2.5cm bins, and function (3) was only evaluated at bins whose centres lay within the boundary. In both environments, Δθ was no smaller than 5.7°.

The best-fitting BVC for each neural rate map was defined by varying the values of *d*, Φ and *σ*_0_ and conducting an exhaustive search that maximised the Pearson’s r correlation between the BVC map and the neural rate map. r_max_ was defined as the correlation between the rate map and the best-fitting BVC map. *d* varied between 0cm and 32.5cm (40cm in circular arenas), in 2.5cm steps, Φ varied between 0° and 354°, in 6° steps, and θ_0_ could take the values 6.2cm, 12.2cm, 20.2cm or 30.2cm. In total the search set consisted of 3120 BVC model maps in square environments (3840 in circular arenas). Following ^25^, the value of θ_*rad*_ was held constant at 0.2 radians, and *θ* at 183cm.

### Assessing statistical significance of BVC fit to neural data

BVC fits to neural rate maps were defined as significant on the basis of a comparison to Pearson’s-r values derived from fitting model maps to spatially shuffled neural data. Spatially-shuffled neural data was produced by shifting the spike train by a random amount between 20 sec, and trial duration minus 20 sec. Rate maps were then constructed using the method described above (“Construction of Firing Rate Maps”). Spatial shuffling was repeated until there was a population of 120,000 shuffled rate maps, for each brain area and age group. Model BVCs were fit to these spatially shuffled rate maps as described above (“Construction of BVC model maps”). For each brain area and age group, the 99^th^ percentile of the Pearson’s-r fit values from spatially shuffled data was taken as the threshold for defining a model fit to (non-shuffled) neural data as significant. An additional threshold, determining a minimum level of spatial specificity for BVCs was also used, which was defined as the 75^th^ percentile of Spatial Information scores (see below, “Assessing spatial tuning and stability of BVCs”) derived from the same population of shuffled data. The 75^th^ percentile was used as the intended purpose of the spatial information threshold was not to reject a null hypothesis of non-spatial firing, but instead set a requirement for a BVC to demonstrate minimal spatial selectivity. The threshold Spatial Information values used were: P16-18, 0.0467; P19-P21, 0.0487; P22-25, 0.0489; Adult, 0.0580.

Neurons were classified as BVCs if they satisfied the above criteria on either of two trials run in the square open arena. The false positive classification rate for each neuron was assumed to be 1.49%, calculated under the assumptions that the chances of satisfying the BVC fit and Spatial Information criteria were independent, and that the chances of satisfying the criteria in either of the two square open field trials were independent. The 95% significance level for the percentage of neurons classified as BVCs at any given age was calculated as the 95^th^ percentile of a binomial distribution based on N samples (where N is the total number of neurons recorded at that age), and a 1.49% success probability.

### Assessing spatial tuning and stability of BVCs

The spatial specificity of neuron firing was assessed using Spatial Information, defined as the mutual information I(R|X) between firing rate R and location X:

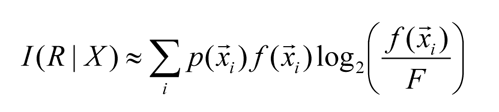

where *p(x_1_*)is the probability for the animal being at location *x_i_*, *f*(*x_i_*) is the firing rate observed at *x_i_*, and F is the overall firing rate of the cell (Skaggs et al., 1996). I(R|X) was then divided by the mean firing rate of the cell, giving an estimate in bits/spike. The spatial stability of BVCs was assessed using the Pearson’s-r correlation of rate maps from temporally adjacent square open field trials. Overall age trend in Spatial Information and Stability were tested using 1-way ANOVA (between subjects factor age), and post-hoc pairwise comparisons using Tukey’s HSD.

### Statistical testing of d, Φ, and σ_0_ distributions

For each BVC, the *d*, Φ, and σ_0_ values were defined as those of the best-fitting model, in the open field trial with the highest fit-maximised Pearson’s-r. Overall age trends in median *d*, and σ_0_ were tested using a Kruskal-Wallis test (between-subjects factor age). To quantify the observed clustering of Φ tunings at the cardinal compass points, a Rayleigh test for unidirectional departure from uniformity was performed on Φ angle data that had first been quadrupled, and then wrapped onto the interval 0-360°. This test is well-suited to detect multimodal departure from uniformity, in cases where a clear hypothesis predicting *n*-fold circular symmetry of the modes exists ^66^. Overall development trends in the proportion of BVCs with Φ tunings oriented to a wall (defined as cardinal compass points ±12°) was tested using a χ^2^ test. Differences in the proportion of wall-oriented Φ between long- and short-distance tuned BVCs (defined as d>10cm and d≤10cm, respectively) were tested using χ^2^ test, post-hoc pairwise comparisons were performed using a z-test for proportions. The distribution of Φ tunings in the circular arena was defined as that of the model that best fit the circular arena neural rate map, and distributions were tested using the Rayleigh test on quadrupled, wrapped data. The mean rotation of BVC ensembles, between the square and circular arenas, was defined as the circular mean of the signed differences in Φ between arenas. BVCs with *d* > 25cm were excluded from this estimate, as their rotations could be ambiguous across a 180° symmetry. The circular dispersion of rotations around ensemble means was assessed using the Kappa test of circular concentration, on the population of the differences between Φ rotation for each BVC, and the respective ensemble mean rotation. To obtain a reliable estimate of mean rotation, only ensembles with ≥5 simultaneously recorded BVCs were included in the analysis: in the following test for 4-fold symmetric clustering at the population level, all developing rats were analysed together, to compensate for the reduced number of BVCs resulting from this restriction. Analysis of changes in 4-fold symmetrical clustering within ensembles used all BVCs and only ensembles with ≥5 simultaneously recorded BVCs.

### BVC responses to inserted barriers

First, for each BVC the appropriate barrier orientation for assessing responses was determined on the basis of the directional tuning: BVCs with Φ tunings 54°≥126° or 234°≥306° (where 0° is East) were defined as north-south tuned BVCs, and responses were assessed using an east-west oriented barrier. BVCs with Φ tunings ≤36°, 144°≥216° or ≥324° were defined as east-west tuned BVCs, and responses were assessed using a north-south oriented barrier. Those BVC whose Φ tunings fell outside these classifications were not considered as unambiguously north-south or east-west oriented, and were excluded from the barrier analysis. A BVC was only included in the analysis if a trial with the appropriate barrier orientation had been conducted in that session, and the *d* tuning of the BVC was ≤15cm. Following this, BVC responses to barriers were assayed using a method similar to that described in ^30^. Briefly, the changes in firing rate were measured in two zones defined relative to barrier position: each zone was as long as the barrier in the dimension parallel to the barrier, and extended from the barrier, to 12.5cm away from the barrier, in the orthogonal dimension. For each BVC, the two zones were designated distal and proximal on the basis of the BVC’s directional tuning, with the distal zone being on the opposite side of the barrier to the direction of Φ. The absolute rate barrier response was then defined as the change in the summed values of rate map bins within each zone, between the barrier trial and the preceding open field square. The normalised response was defined as the absolute response, divided by the sum of the summed rates in the barrier and open field trials. Overall age trends in barrier response were tested using a mixed design ANOVA, including age as a between-subjects factor and zone (distal versus proximal) as a within-subjects factor. Post-hoc pairwise comparisons were conducted using Simple Main Effects.

### Comparison of subiculum and mEC neural data

The mEC dataset analysed here was previously described in ^30^: full details of data recording conditions are contained therein. Data was shared in the form of position tracking records and spike times of isolated single units (the same set as described in the original study). All further analysis, including construction of rate maps, fitting of BVCs and determining fit significance using shuffled data, was performed as described for subiculum data, above. Neurons were defined as border cells using the same method as ^30^, namely, both the Border Score ^23^ and the Spatial Information (see above) of a neuron’s rate map needed to exceed the 95^th^ percentiles of populations of Border and Spatial Information scores derived from age-matched, spatially-shuffled data. The Spatial Information, stability and BVC tuning properties of both BVCs and Border Cells identified in the mEC were analysed as for subiculum BVCs, as described above.

---

**Supplemental Figure 1.**
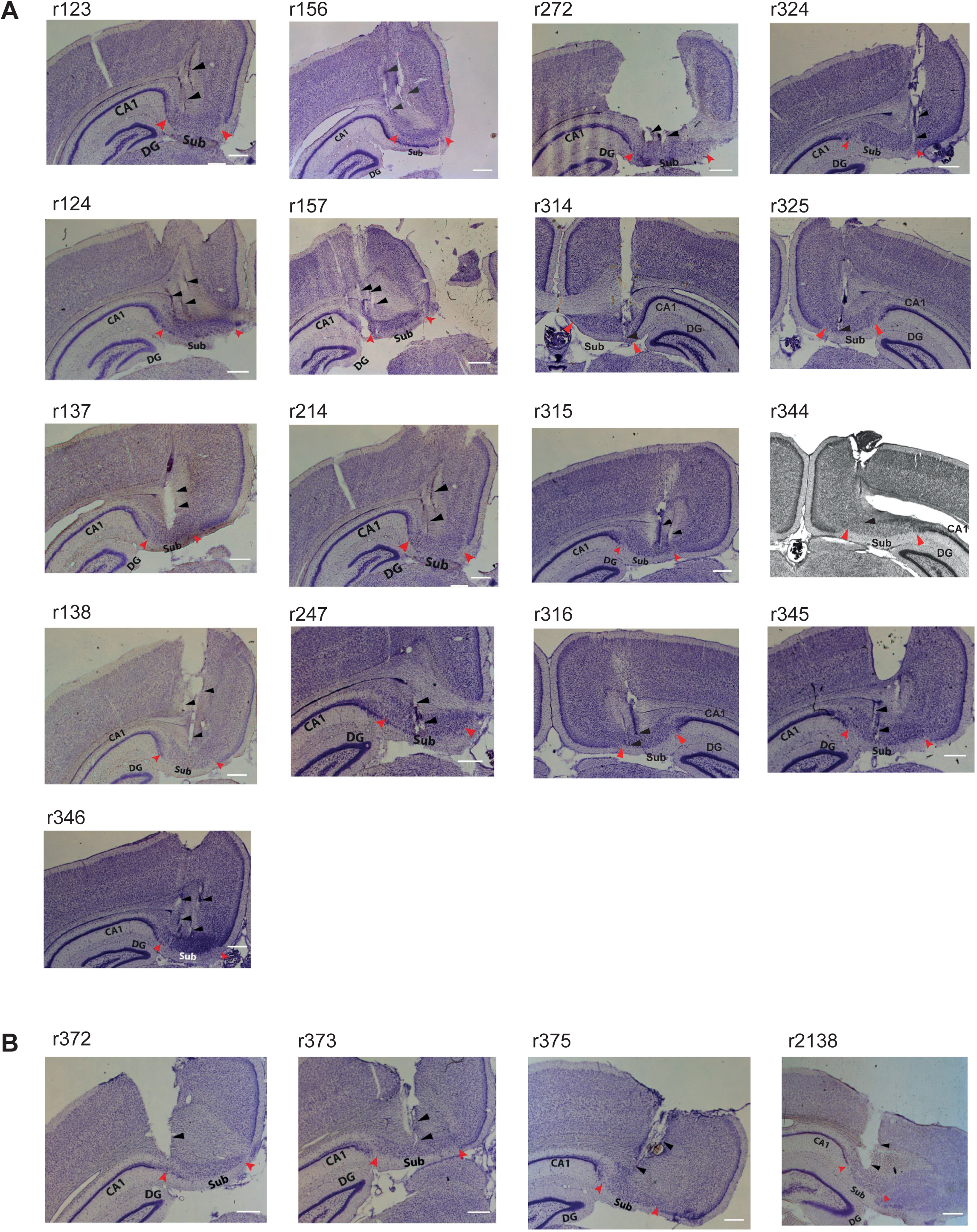
Representative post-mortem histological sections (one per rat), showing tetrode positions in the subiculum, for developing (**A**) and adult (**B**) subjects. Red arrowheads show the extent of the subiculum, black arrowheads indicate tetrode positions. CA1, Hippocampus Cornu Ammonis field 1, DG, Dentate Gyrus, Sub, Subiculum. Scale bar shows 500μm

**Supplemental Figure 2.**
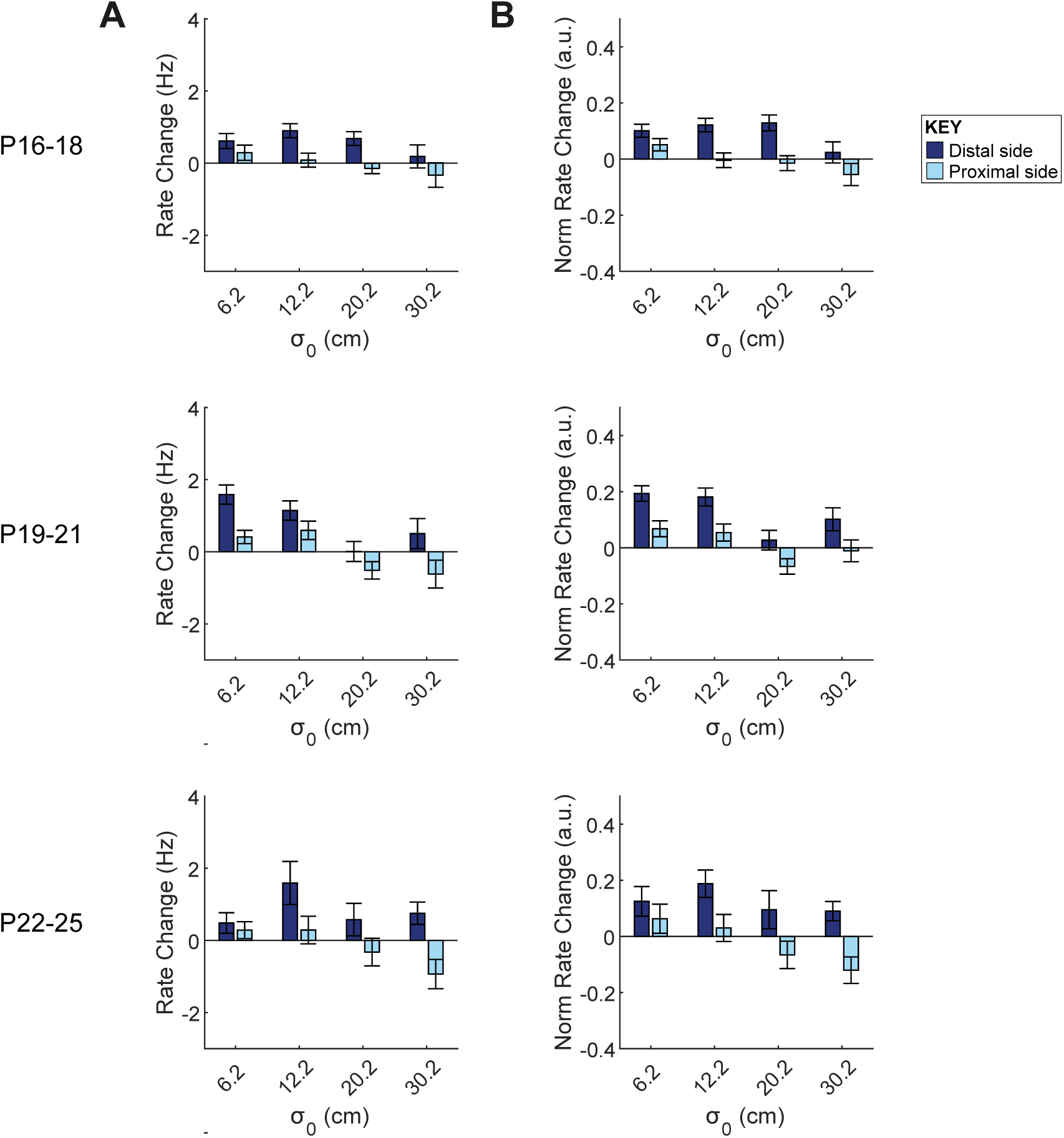
BVC responses to inserted barriers in developing rats, split by σ_0_ of best-fit model in the base- line trial. Broader tuned BVCs showed a trend towards inhibited firing on the proximal side of the barrier, as develop- ment proceeded, but this trend did not reach significance. **(A)** Mean absolute barrier rate change score (±SEM) for BVCs in each age group. Dark blue bars show rate changes on distal side of the barrier, light blue bars on the proximal side of the barrier, bar pairs show BVC responses for each σ_0_ value. There is a significant effect of σ_0_ on proximal side firing only at P19-21, though the trend approaches signifi- cance at P22-25 (ANOVA, factor σ_0_, P16-18, F_(3,218)_=1.13; p=0.33; P19-21, F_(3,159)_=5.06; p=0.002; P22-25, F_(3,81)_=2.65; p=0.055). At P19-P21, no individual σ_0_ groups were significantly different from zero, following a Sidak correction for 4 comparisons (corrected α=0.0127; σ_0_=6.2, p=0.015; σ_0_=12.2, p=0.014; σ_0_=20.2, p=0.022; σ_0_=30.2, p=0.060). Likewise, notwithstanding the lack of an overall effect of baseline best-fit σ at P22-P25, no individual σ_0_ groups were significantly different from zero (corrected α=0.0127; σ_0_=6.2, p=0.88; σ_0_=12.2, p=0.75; σ_0_=20.2, p=0.20; σ_0_=30.2, p=0.018). **(B)** Mean normalised barrier rate change score (±SEM) for BVCs in each age group. There is a significant effect of σ_0_ on proximal side firing only at P19-21, though the trend approaches significance at P16-18 and P22-25 (ANOVA, factor σ_0_, P16-18, F_(3,218)_= 2.48; p=0.061; P19-21, F_(3,159)_= 3.00; p=0.032; P22-25, F_(3,81)_=2.66; p=0.053). At P19-P21, the σ_0_ =20.2 group was significantly different from zero, following a Sidak correction for 4 comparisons (corrected α=0.0127; σ_0_=6.2, p=0.023; σ_0_=12.2, p=0.042; σ_0_=20.2, p=0.012; σ_0_=30.2, p=0.39). Likewise, notwithstanding the lack of an over- all effect of baseline best-fit σ at P22-P25, the σ_0_=30.2 group was significantly different from zero (corrected α=0.0127; σ_0_=6.2, p=0.88; σ_0_=12.2, p=0.73; σ_0_=20.2, p=0.099; σ_0_=30.2, p=0.011).

**Supplemental Figure 3.**
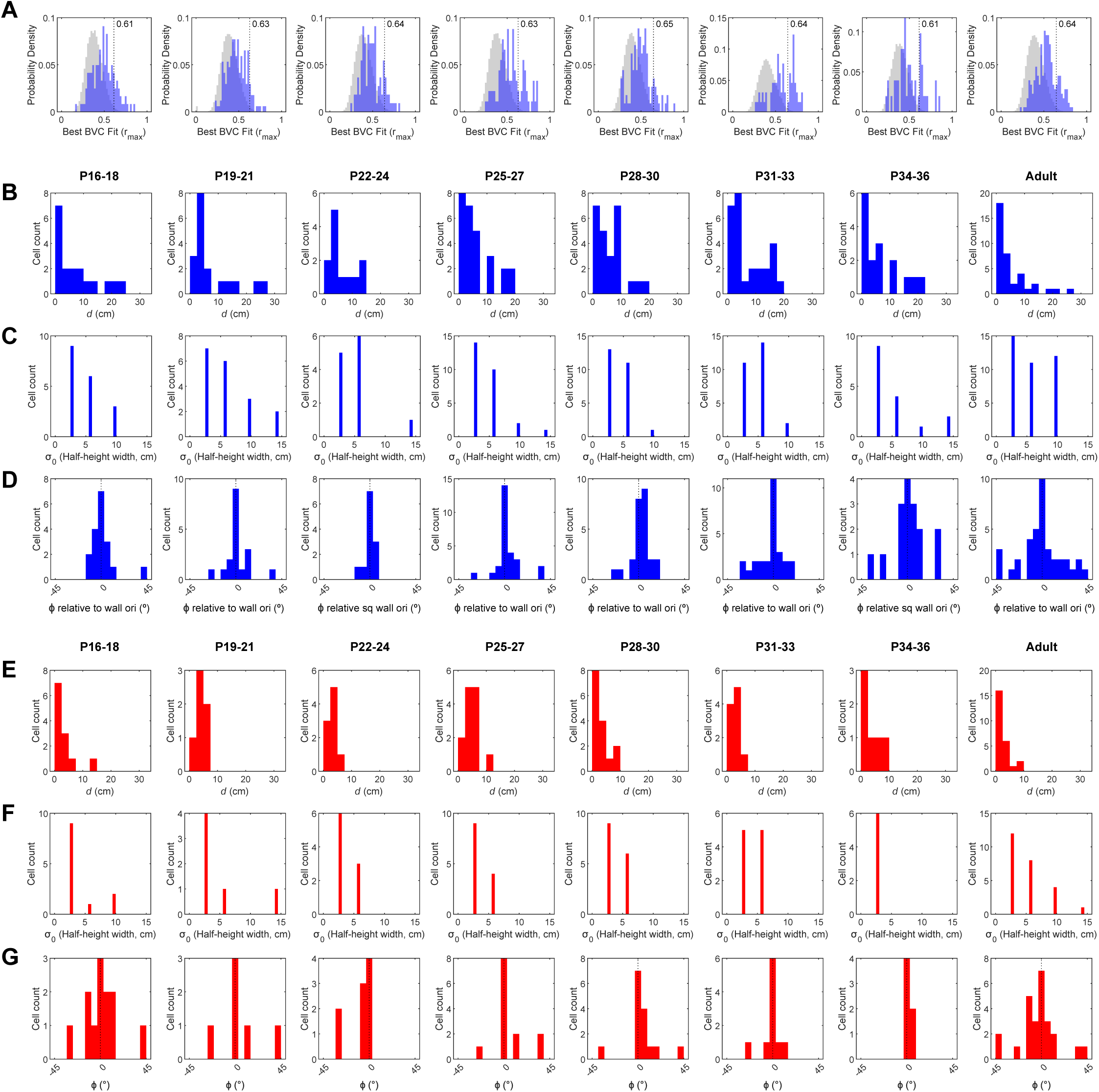
Extended characterisation of BVCs and Border Cells in mEC. (**A**) Distributions of correlations between neuronal firing rate maps and the best-fitting BVC map (r_max_), for mEC data. Blue histograms show mEC data r_max_, grey histograms show r_max_ based on shuffled data. Each histogram axes refers to an age group. Vertical dashed lines show the threshold r r_max_ distributions, within each age group. Max for BVC classification, defined as the 99^th^ percentile of the spike-shifted (**B**-**D**) Histograms of receptive field tuning properties for mEC BVCs in each age group: (**B**) Histograms showing all mEC BVC *d* tunings in each age group. (**C**) Counts of mEC BVC σ_0_ tunings in each age group. σ_0_ tunings are expressed as the half-height width of a BVC field with given σ_0_, assuming *d*=0 and φ=0. (**D**) Histo- grams of all mEC BVC φ tunings mapped onto one 90° quadrant. 0° indicates φ tunings aligned to a wall. (**E**-**G**) Histograms of receptive field tuning properties for mEC Border Cells in each age group: (**E**) Histograms showing all mEC Border Cell *d* tunings in each age group. (**F**) Counts of mEC Border Cell σ_0_ tunings in each age group. σ_0_ tunings are expressed as the half-height width of a BVC field with given σ_0_, assuming *d*=0 and φ=0. (**G**) Histograms of all mEC Border Cell φ tunings mapped onto one 90° quadrant. 0° indicates φ tunings aligned to a wall. Receptive field properties of border cells were derived from the best-fitting model BVC, irrespective of whether the model BVC was determined to be an above-chance fit for the neural firing.

## References

1. O’Keefe, J. & Dostrovsky, J. The hippocampus as a spatial map. Preliminary evidence from unit activity in the freely-moving rat. Brain Res. 34, 171–175 (1971).

2. Taube, J. S., Muller, R. U. & Ranck Jr., J. B. Head-direction cells recorded from the postsubiculum in freely moving rats. I. Description and quantitative analysis. J.Neurosci. 10, 420–435 (1990).

3. Hafting, T., Fyhn, M., Molden, S., Moser, M. B. & Moser, E. I. Microstructure of a spatial map in the entorhinal cortex. Nature 436, 801–806 (2005).

4. O’Keefe, J. & Nadel, L. The hippocampus as a cognitive map. (Oxford University Press, 1978).

5. Poulter, S., Hartley, T. & Lever, C. The Neurobiology of Mammalian Navigation. Curr Biol 28, R1023–R1042 (2018).

6. Muller, R. U. & Kubie, J. L. The effects of changes in the environment on the spatial firing of hippocampal complex-spike cells. J.Neurosci. 7, 1951–1968 (1987).

7. Gothard, K. M., Skaggs, W. E. & McNaughton, B. L. Dynamics of mismatch correction in the hippocampal ensemble code for space: interaction between path integration and environmental cues. J.Neurosci. 16, 8027–8040 (1996).

8. Chen, G., King, J. A., Burgess, N. & O’Keefe, J. How vision and movement combine in the hippocampal place code. Proc Natl Acad Sci U S A 110, 378–83 (2013).

9. Stewart, S., Jeewajee, A., Wills, T. J., Burgess, N. & Lever, C. Boundary coding in the rat subiculum. Philos Trans R Soc Lond B Biol Sci 369, 20120514 (2014).

10. Barry, C. et al. The boundary vector cell model of place cell firing and spatial memory. Rev.Neurosci. 17, 71–97 (2006).

11. McNaughton, B. L. et al. Deciphering the hippocampal polyglot: the hippocampus as a path integration system. J.Exp.Biol. 199 (Pt 1, 173–185 (1996).

12. Touretzky, D. S. & Redish, A. D. Theory of rodent navigation based on interacting representations of space. Hippocampus 6, 247–270 (1996).

13. Hasselmo, M. E. Grid cell mechanisms and function: contributions of entorhinal persistent spiking and phase resetting. Hippocampus 18, 1213–1229 (2008).

14. Burgess, N. Grid cells and theta as oscillatory interference: theory and predictions. Hippocampus 18, 1157–1174 (2008).

15. Burak, Y. & Fiete, I. R. Accurate path integration in continuous attractor network models of grid cells. PLoS.Comput.Biol. 5, e1000291 (2009).

16. Hardcastle, K., Ganguli, S. & Giocomo, L. M. Environmental boundaries as an error correction mechanism for grid cells. Neuron 86, 827–39 (2015).

17. Cheng, K. A purely geometric module in the rat’s spatial representation. Cognition 23, 149–178 (1986).

18. Hermer, L. & Spelke, E. S. A geometric process for spatial reorientation in young children. Nature 370, 57–59 (1994).

19. Cheng, K., Huttenlocher, J. & Newcombe, N. S. 25 years of research on the use of geometry in spatial reorientation: a current theoretical perspective. Psychon Bull Rev 20, 1033–1054 (2013).

20. Doeller, C. F., King, J. A. & Burgess, N. Parallel striatal and hippocampal systems for landmarks and boundaries in spatial memory. Proc Natl Acad Sci U S A 105, 5915– 5920 (2008).

21. Lee, S. A. et al. Electrophysiological Signatures of Spatial Boundaries in the Human Subiculum. J Neurosci 38, 3265–3272 (2018).

22. Hartley, T., Trinkler, I. & Burgess, N. Geometric determinants of human spatial memory. Cognition 94, 39–75 (2004).

23. Solstad, T., Boccara, C. N., Kropff, E., Moser, M. B. & Moser, E. I. Representation of geometric borders in the entorhinal cortex. Science (1979) 322, 1865–1868 (2008).

24. Savelli, F., Yoganarasimha, D. & Knierim, J. J. Influence of boundary removal on the spatial representations of the medial entorhinal cortex. Hippocampus 18, 1270–82 (2008).

25. Hartley, T., Burgess, N., Lever, C., Cacucci, F. & O’Keefe, J. Modeling place fields in terms of the cortical inputs to the hippocampus. Hippocampus 10, 369–379 (2000).

26. Lever, C., Burton, S., Jeewajee, A., O’Keefe, J. & Burgess, N. Boundary vector cells in the subiculum of the hippocampal formation. J.Neurosci. 29, 9771–9777 (2009).

27. Cohen, L., Vinepinsky, E., Donchin, O. & Segev, R. Boundary vector cells in the goldfish central telencephalon encode spatial information. PLoS Biol 21, e3001747 (2023).

28. Stangl, M. et al. Boundary-anchored neural mechanisms of location-encoding for self and others. Nature 589, 420–425 (2021).

29. Muessig, L., Hauser, J., Wills, T. J. & Cacucci, F. A Developmental Switch in Place Cell Accuracy Coincides with Grid Cell Maturation. Neuron 86, 1167–1173 (2015).

30. Bjerknes, T. L., Moser, E. I. & Moser, M.-B. Representation of geometric borders in the developing rat. Neuron 82, 71–8 (2014).

31. Kim, S. M., Ganguli, S. & Frank, L. M. Spatial Information Outflow from the Hippocampal Circuit: Distributed Spatial Coding and Phase Precession in the Subiculum. Journal of Neuroscience 32, 11539–11558 (2012).

32. Ledergerber, D. et al. Task-dependent mixed selectivity in the subiculum. Cell Rep 35, 109175 (2021).

33. Olson, J. M., Tongprasearth, K. & Nitz, D. A. Subiculum neurons map the current axis of travel. Nat Neurosci 20, 170–172 (2017).

34. Poulter, S., Lee, S. A., Dachtler, J., Wills, T. J. & Lever, C. Vector trace cells in the subiculum of the hippocampal formation. Nature Neuroscience 2020 24:2 24, 266–275 (2020).

35. Kitanishi, T., Umaba, R. & Mizuseki, K. Robust information routing by dorsal subiculum neurons. Sci Adv 7, (2021).

36. Aggleton, J. P. & Christiansen, K. The subiculum: the heart of the extended hippocampal system. Prog Brain Res 219, 65–82 (2015).

37. Donato, F., Jacobsen, R. I., Moser, M.-B. & Moser, E. I. Stellate cells drive maturation of the entorhinal-hippocampal circuit. Science (1979) 355, eaai8178 (2017).

38. Høydal, Ø. A., Skytøen, E. R., Andersson, S. O., Moser, M.-B. & Moser, E. I. Object- vector coding in the medial entorhinal cortex. Nature 568, 400–404 (2019).

39. Deshmukh, S. S. & Knierim, J. J. Influence of local objects on hippocampal representations: Landmark vectors and memory. Hippocampus 23, 253–267 (2013).

40. Bicanski, A. & Burgess, N. Neuronal vector coding in spatial cognition. Nat Rev Neurosci 21, 453–470 (2020).

41. O’Keefe, J. & Burgess, N. Geometric determinants of the place fields of hippocampal neurons. Nature 381, 425–428 (1996).

42. Kosaki, Y., Austen, J. M. & McGregor, A. Overshadowing of geometry learning by discrete landmarks in the water maze: effects of relative salience and relative validity of competing cues. J Exp Psychol Anim Behav Process 39, 126–39 (2013).

43. Twyman, A. D., Holden, M. P. & Newcombe, N. S. First Direct Evidence of Cue Integration in Reorientation: A New Paradigm. Cogn Sci 42 Suppl 3, 923–936 (2018).

44. Stensola, T., Stensola, H., Moser, M. B. & Moser, E. I. Shearing-induced asymmetry in entorhinal grid cells. Nature 518, 207–212 (2015).

45. Krupic, J., Bauza, M., Burton, S., Barry, C. & O’Keefe, J. Grid cell symmetry is shaped by environmental geometry. Nature 518, 232–235 (2015).

46. Wills, T. J., Cacucci, F., Burgess, N. & O’Keefe, J. Development of the hippocampal cognitive map in preweanling rats. Science (1979) 328, 1573–1576 (2010).

47. Langston, R. F. et al. Development of the spatial representation system in the rat. Science (1979) 328, 1576–1580 (2010).

48. Alexander, A. S. et al. Egocentric boundary vector tuning of the retrosplenial cortex. Sci Adv 6, (2020).

49. Bellmund, J. L. S. et al. Deforming the metric of cognitive maps distorts memory. Nat Hum Behav 4, 177–188 (2020).

50. Bye, C. M., Hong, N. S., Moore, K., Deibel, S. H. & McDonald, R. J. The effects of pool shape manipulations on rat spatial memory acquired in the Morris water maze. Learn Behav 47, 29–37 (2019).

51. Pearce, J. M., Good, M. A., Jones, P. M. & McGregor, A. Transfer of spatial behavior between different environments: implications for theories of spatial learning and for the role of the hippocampus in spatial learning. J.Exp.Psychol.Anim Behav.Process 30, 135–147 (2004).

52. McGregor, A., Hayward, A. J., Pearce, J. M. & Good, M. A. Hippocampal lesions disrupt navigation based on the shape of the environment. Behav.Neurosci. 118, 1011–1021 (2004).

53. Rathore, S. The sensory and behavioural determinants of neuronal activity in the Subiculum (PhD Thesis). (UCL (University College London), 2017).

54. Zhang, K. Representation of spatial orientation by the intrinsic dynamics of the head- direction cell ensemble: a theory. J.Neurosci. 16, 2112–2126 (1996).

55. Redish, A. D., Elga, A. N. & Touretzky, D. S. A coupled attractor model of the rodent head direction system. Network 7, 671 (1996).

56. Bicanski, A. & Burgess, N. A neural-level model of spatial memory and imagery. Elife 7, (2018).

57. Sharma, A., Nair, I. R. & Yoganarasimha, D. Attractor-like Dynamics in the Subicular Complex. J Neurosci 42, 7594–7614 (2022).

58. Wills, T. J., Muessig, L. & Cacucci, F. The development of spatial behaviour and the hippocampal neural representation of space. Philos Trans R Soc Lond B Biol Sci 369, 20130409 (2014).

59. Scott, R. C., Richard, G. R., Holmes, G. L. & Lenck-Santini, P. P. Maturational dynamics of hippocampal place cells in immature rats. Hippocampus (2010).

60. Bullens, J. et al. The role of landmarks and boundaries in the development of spatial memory. Dev.Sci. 13, 170–180 (2010).

61. Glöckner, F., Schuck, N. W. & Li, S. C. Differential prioritization of intramaze cue and boundary information during spatial navigation across the human lifespan. Sci Rep 11, (2021).

62. Lehnung, M. et al. Development of spatial memory and spatial orientation in preschoolers and primary school children. British Journal of Psychology 89, 463–480 (1998).

63. Overman, W. H., Pate, B. J., Moore, K. & Peuster, A. Ontogeny of place learning in children as measured in the Radial Arm Maze, Morris Search Task, and Open Field Task. Behavioral Neuroscience 110, 1205–1228 (1996).

64. Twyman, A. D., Newcombe, N. S. & Gould, T. J. Malleability in the development of spatial reorientation. Dev Psychobiol 55, 243–255 (2013).

65. Chiandetti, C. & Vallortigara, G. Is there an innate geometric module? Effects of experience with angular geometric cues on spatial re-orientation based on the shape of the environment. Anim Cogn 11, 139–146 (2008).

66. Landler, L., Ruxton, G. D. & Malkemper, E. P. Circular data in biology: advice for effectively implementing statistical procedures. Behav Ecol Sociobiol 72, (2018).

